# Measuring G Protein Activation with Spectrally Resolved Fluorescence Fluctuation Spectroscopy

**DOI:** 10.1101/2021.11.03.467169

**Authors:** Daniel J. Foust, David W. Piston

## Abstract

G protein-coupled receptor signaling has been posited to occur through either collision coupling or pre-assembled complexes with G protein transducers. To investigate the dynamics of G protein signaling, we introduce fluorescence covariance matrix analysis (FCMA), a novel implementation of fluorescence cumulant analysis applied to spectrally resolved fluorescence images. We labeled the GPCR, Gα, and Gβγ units with distinct fluorescent protein labels and we applied FCMA to measure directly the complex formation during stimulation of dopamine and adrenergic receptors. To determine the prevalence of hetero-oligomers, we compared the GPCR data to those from control samples expressing three fluorescent protein labels with known stoichiometries. Interactions between Gα and Gβγ subunits determined by FCMA were sensitive to stimulation with GPCR ligands. However, GPCR/G protein interactions were too weak to be distinguished from background. These findings support a collision coupling mechanism rather than pre-assembled complexes for the two GPCRs studied.

## Introduction

Fluorescence fluctuation spectroscopy (FFS) is a set of statistical techniques used to extract physical parameters from fluorescence signals by using physical models of fluorescence detection (1). In biological applications, the most frequently measured parameters are diffusion coefficients, concentrations, and the molecular brightness of fluorescently labeled biomolecules (2–4). Imaging-FFS, by analyzing multidimensional fluorescence images, has become increasingly popular due to its potential to provide spatially resolved information (5–8). FFS is also used to analyze samples containing multiple chromophores and provide information about heteromeric molecular interactions (9, 10). Towards measuring an increasing number of molecular components, multicolor FFS has been expanded to utilize spectrally resolved detection. In these systems, a prism or diffraction grating is used to redirect photons onto an array of detectors so that the energy of incident photoelectrons is known more precisely than in systems based on dichroic mirrors and multiple independent detectors (11–13). Recent developments have paired spectral detection with spectral unmixing techniques for better signal-to-noise ratios (SNRs) (14–16), and as many as four chromophores have been used simultaneously in live cell experiments (17).

G protein coupled receptors (GPCRs) are physiologically vital cell surface receptors (18). Their downstream effects are predominantly mediated by G proteins (19–21). Trimeric G proteins are heterocomplexes composed of α, β, and γ subunits (22). Together, the β and γ subunits, form a single functional signaling unit and are not active when unpaired. Canonically, GPCRs catalyze the dissociation of Gαβγ into active Gα and Gβγ signaling units that can participate in downstream signaling events. The interactions between GPCRs and G proteins have been probed extensively using several fluorescence-based techniques including resonance energy transfer, bimolecular complementation, single molecule tracking, polarization, and FFS (23, 24).

An outstanding issue in GPCR signaling is the degree to which receptors and G proteins form stable complexes (25). Collision coupling models posit that GPCRs and G proteins move independently in the plasma membrane in the absence of stimulating ligands and are limited to transient interactions (26–28). Pre-assembly models assert that GPCRs and G proteins interact constitutively in stable complexes (29–31). Kinetic models have been developed to accommodate both signaling mechanisms (32), but experimental evidence to date has been contradictory regarding which mechanism is dominant (24). Multicolor FFS approaches are well-suited to address this problem because they enable us to observe the principal components of GPCR signal transduction simultaneously and in real time.

To investigate the mechanisms underlying GPCR activation, we introduce fluorescence covariance matrix analysis (FCMA), which is a new approach to analyzing spectrally resolved imaging data that provides detection and quantification of multi-chromic complexes. Additionally, FCMA provides a simple graphical interpretation of complex formation. FCMA is an extension of cumulant-based approaches applied to spectrally resolved imaging data (8, 33–35). We applied FCMA to 23 channel images of tri-chromic samples expressing permutations of green, yellow, and red fluorescent protein monomers and dimers expressed on the plasma membrane. Using FCMA, we can detect and measure trimeric interactions without invoking higher order correlations (17).

We applied FCMA to the analysis of G protein signaling mechanisms for two Gαi-coupled GPCRs, DRD2 and ADRA2A, whose exogenous ligands are dopamine and epinephrine, respectively. The Gαi subset of G proteins are involved in the inhibition of cAMP production through interactions with adenylyl cyclase (36). In co-expression experiments with labeled Gαi and Gβγ functional signaling units, we monitored these three components during the signaling event. The Gα/Gβγ interactions were sensitive to stimulation with GPCR agonists. However, we did not observe ternary interactions between GPCRs, Gαi, and Gβγ units, which is consistent with the absence of pre-assembled complexes.

## Results

### Fluorescence Covariance Matrix Analysis of CD86 Controls

To calibrate FCMA as a tool for analyzing samples expressing multiple chromophores, we used the monomeric plasma membrane protein CD86 as a scaffold for permutations of fluorescent heteromers. Three fluorescent proteins were used, mEGFP (G), mEYFP (Y), and mCherry2 (R). Three fluorescent monomers were analyzed, CD86-G, CD86-Y, and CD86-R, as well as five monomer/heteromer combinations, CD86-G + CD86-Y + CD86-R (three monomers), R-CD86-Y-G (trimer), CD86-Y-G + CD86-R (GY dimer with R monomer), CD86-G + R-CD86-Y (G monomer with YR dimer), and R-CD86-G + CD86-Y (GR dimer with Y monomer) (Fig. 1). These combinations were expressed in HEK 293 cells and the plasma membrane adjacent to the cover glass was imaged using confocal microscopy.

**FIGURE 1.**
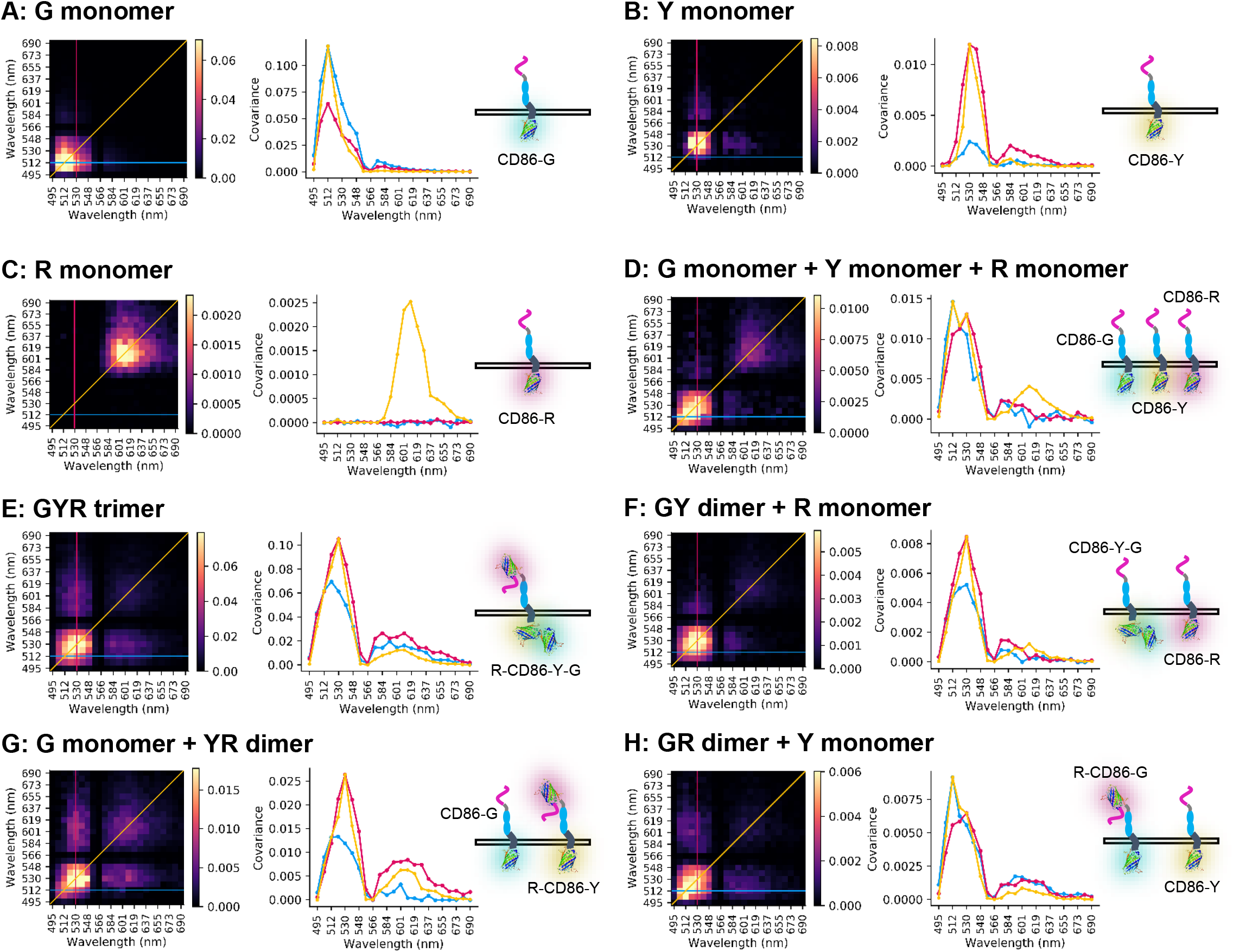
Representative covariance matrices 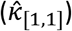 for cells expressing fluorescent monomers and heteromers in several permutations. A) CD86-G expressed alone. B) CD86-Y. C) CD86-R. D) Triple transfection of CD86-G, CD86-Y, and CD86-R. E) Fluorescent trimer, R-CD86-Y-G. F) Cotransfection of fluorescent heterodimer CD86-Y-G and fluorescent monomer CD86-R. G) Cotransfection of CD86-G and R-CD86-Y. H) Cotransfection of R-CD86-G and CD86-Y. Covariance matrices are on the left of each panel. Middle plots display profiles indicated by lines on covariance matrices. The yellow lines denote the main diagonal. Blue lines denote the line where ordinate is fixed at 512 nm (peak mEGFP detection). The magenta lines denote where the abscissa is fixed at 530 nm (peak mEYFP(Q69K) detection). On the right are cartoon representations of the molecular constructs expressed in each sample.

Samples containing a single chromophore feature a single peak on the main diagonal of the covariance matrix (Fig. 1 A-C). The location of this peak matches the position of the maximum signal in the detection spectrum (Fig. S1 C). When noninteracting species are present in a sample their contributions to the covariance matrix are additive (Fig. 1 D, F-H). When two chromophores interact and act as a single species this has a multiplicative effect on their contribution to the covariance matrix. Interacting chromophores with high spectral overlap produce a broadened peak, as in the case for G and Y (Fig. 1 E-F). If interacting chromophores are well separated spectrally, their contribution to the covariance matrix produces lobes away from the main axis, as is the case for GR and YR interactions (Fig. 1 E, G-H). In the case of ternary interactions, there is both the more prominent, broad peak from the highly overlapping GY interactions and broader off axis lobes from concomitant GR and YR interactions (Fig. 1 E).

### Distribution of Fluorescent Oligomer States by Fluorescence Covariance Matrix Analysis

The covariance matrices and corresponding detection spectra from images of cells expressing three chromophores on CD86 were fit using a seven-component model accounting for the following species: G, Y, R, GY, GR, YR, and GYR. For each component, an apparent number density is determined and the fractional distribution of each chromophore across different oligomer states is found (Fig. 2). The sum of fractional densities for each chromophore is unity. For the triple expression of monomeric proteins, CD86-G, CD86-Y, and CD86-R the largest fractions for each chromophore are observed in monomeric states (Fig. 2 A). We did observe some apparent GY and YR interactions which we take as the background levels for further experiments. GYR fractions were negligible.

**FIGURE 2.**
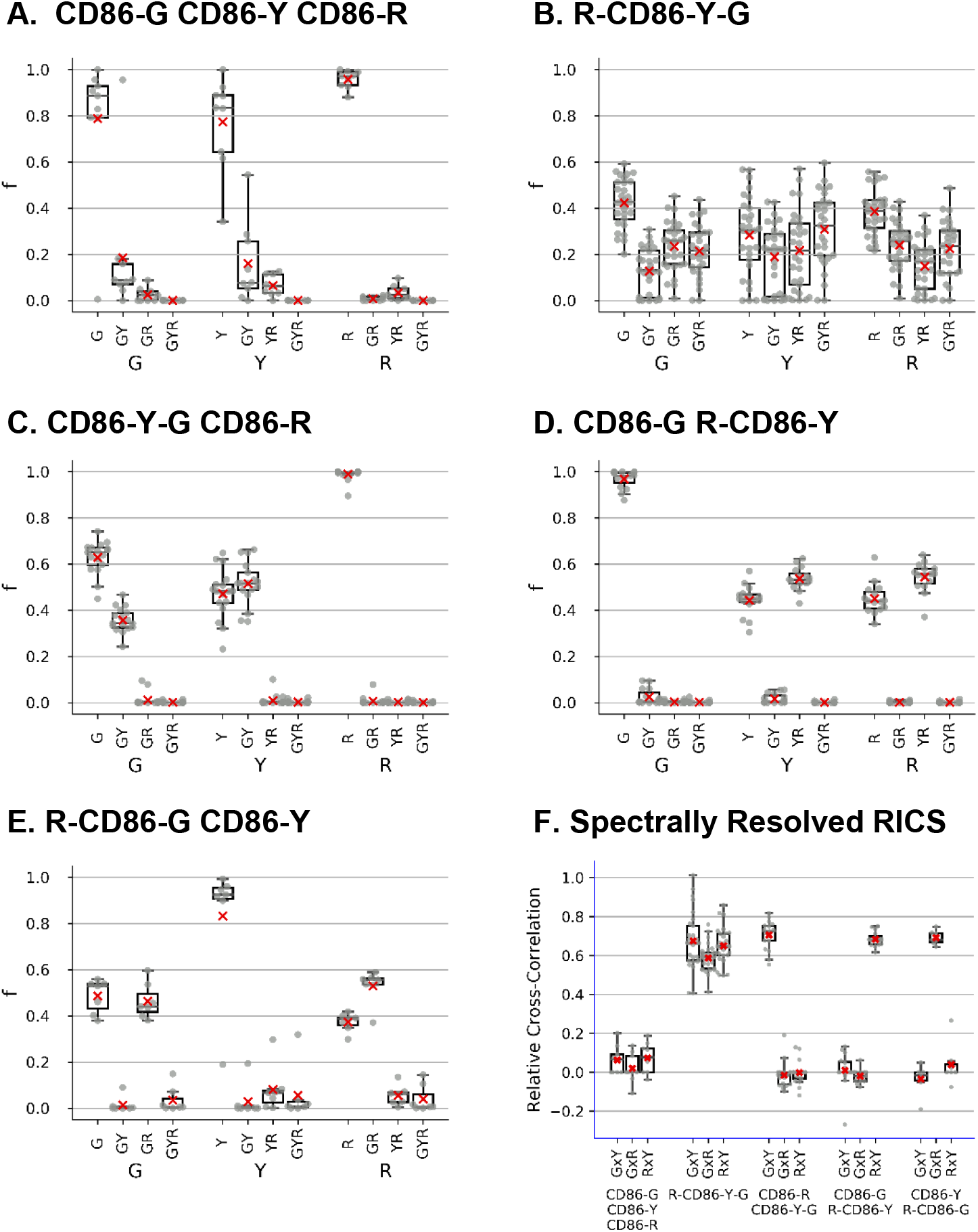
Fractional distribution of chromophores among oligomer states determined by fluorescent covariance matrix analysis. Data is from images of HEK 293 cells expressing fluorescent monomers and heteromers. (A) Cotransfection of CD86-G, CD86-Y, and CD86-R. (B) R-CD86-Y-G expressed alone. (C) Cotransfection of CD86-Y-G and CD86-R. (D) Cotransfection of CD86-G and R-CD86-Y. (E) Cotransfection of R-CD86-G and CD86-Y. (F) Relative cross-correlations determined by spectrally resolved RICS for all permutations.

In cells where the heterotrimer, R-CD86-Y-G was expressed, all three chromophores are found to be spread across several oligomer states (Fig. 2 B). This phenomenon can be understood by considering the existence of dark state fractions for each chromophore. A single protein will only contribute to the GYR (trimer) fraction if all of its constituent chromophores are correctly folded into the fluorescent state. Due to this effect, only small fractions (^~^20-30% per chromophore) appear in the trimeric state. Correspondingly, the dimeric states are populated by contributions from proteins with one chromophore in a dark state and the monomeric states are populated by contributions from proteins with two chromophores in dark states.

Data from cotransfections of dimer and monomer constructs produce similar results. For CD86-Y-G and CD86-R cotransfections, G is distributed predominantly between G (^~^65%) and GY (^~^35%) states (Fig. 2 C). Similarly, Y is split between Y (^~^50%) and GY (^~^50%) states. R is found almost exclusively in its monomeric state. For CD86-G expressed with R-CD86-Y, YR interaction is detected along with smaller fractions in Y and R monomeric states (Fig. 2 D). G is found almost exclusively in its monomeric state. For R-CD86-G expressed with CD86-Y, G and R are split between their respective monomeric states and GR interacting state, whereas Y almost exclusively found in its monomeric state (Fig. 2 E).

### Control Analysis by Spectrally Resolved RICS

To compare with an established method, we processed the same datasets used for FCMA with spectrally resolved raster image correlation spectroscopy (RICS). In contrast with FCMA, the 23 channel spectrally resolved images must be unmixed prior to analysis into three single-color images, each corresponding to a single chromophore (Fig. S1 B-D). Fitting model spatial auto- and cross-correlation functions allows for the calculation of relative cross-correlation amplitudes, a readout of the interaction between pairs of chromophores. Results from RICS analysis (Fig. 2 F) are in good agreement with those found with FCMA. The data show very little interaction between chromophores in triple transfections of G, Y, and R monomers. Images of the R-CD86-Y-G trimer exhibit strong, but not ideal, relative cross-correlations among the three possible pairings. For monomer/dimer coexpressions, CD86-Y-G with CD86-R displays strong GY interactions, R-CD86-Y with CD86-G displays strong YR interaction, and R-CD86-G with CD86-Y displays strong GR interactions (Fig. 2 F). Conventional RICS analysis is limited to concomitant measurements of binary interactions while ternary interactions cannot be detected directly and must be inferred. In contrast, ternary interactions can be quantified directly from covariance matrices (Fig. 1 E, Fig. 2 B).

### Measuring Activation Ligand Induced G Protein Dissociation through DRD2 and ADRA2A

To express all three G proteins at physiologically appropriate and experimentally pragmatic levels, we modified a polycistronic construct introduced by Unen et al (37) to carry mCherry2 tags on GNB1 and GNG2 and an mEYFP(Q69K) tag on GNAI1 (Fig. S2). With this configuration, we labeled the two functional G protein components with Y and R. This construct was expressed in tandem with a G labeled GPCR so that all three components of the GPCR signaling triplet could be monitored simultaneously. The two Gαi-coupled GPCRs studied here were DRD2 (dopamine) and ADRA2A (epinephrine).

GNAI1-Y is primarily a fluorescent monomer, as determined by FCMA, with ^~^50-70% of Y chromophores appearing monomeric before the addition of a GPCR stimulating ligand (Fig. 3). The weak interactions detected between GNAI1-Y and GPCRs G-DRD2 (Fig. 3 A-B) and G-ADRA2A (Fig. 3 C-D) are comparable to background interaction levels found in control experiments featuring CD86 (Fig. 2 A). There is significant interaction between GNAI1-Y and R-GNB1/R-GNG2 with ^~^20% of Y chromophores being found in these interactions. The Y and YR fractions are sensitive to GPCR stimulation by its native ligand. When expressed with G-DRD2, the fraction of Y participating in YR interactions decreases from ^~^20% to ^~^10% after stimulation with 100 μM dopamine (Fig. 3 B). Similarly, when expressed with G-ADRA2A, the YR fraction decreases from ^~^20% to ^~^10% after stimulation with 30 μM epinephrine (Fig. 3 D).

**FIGURE 3.**
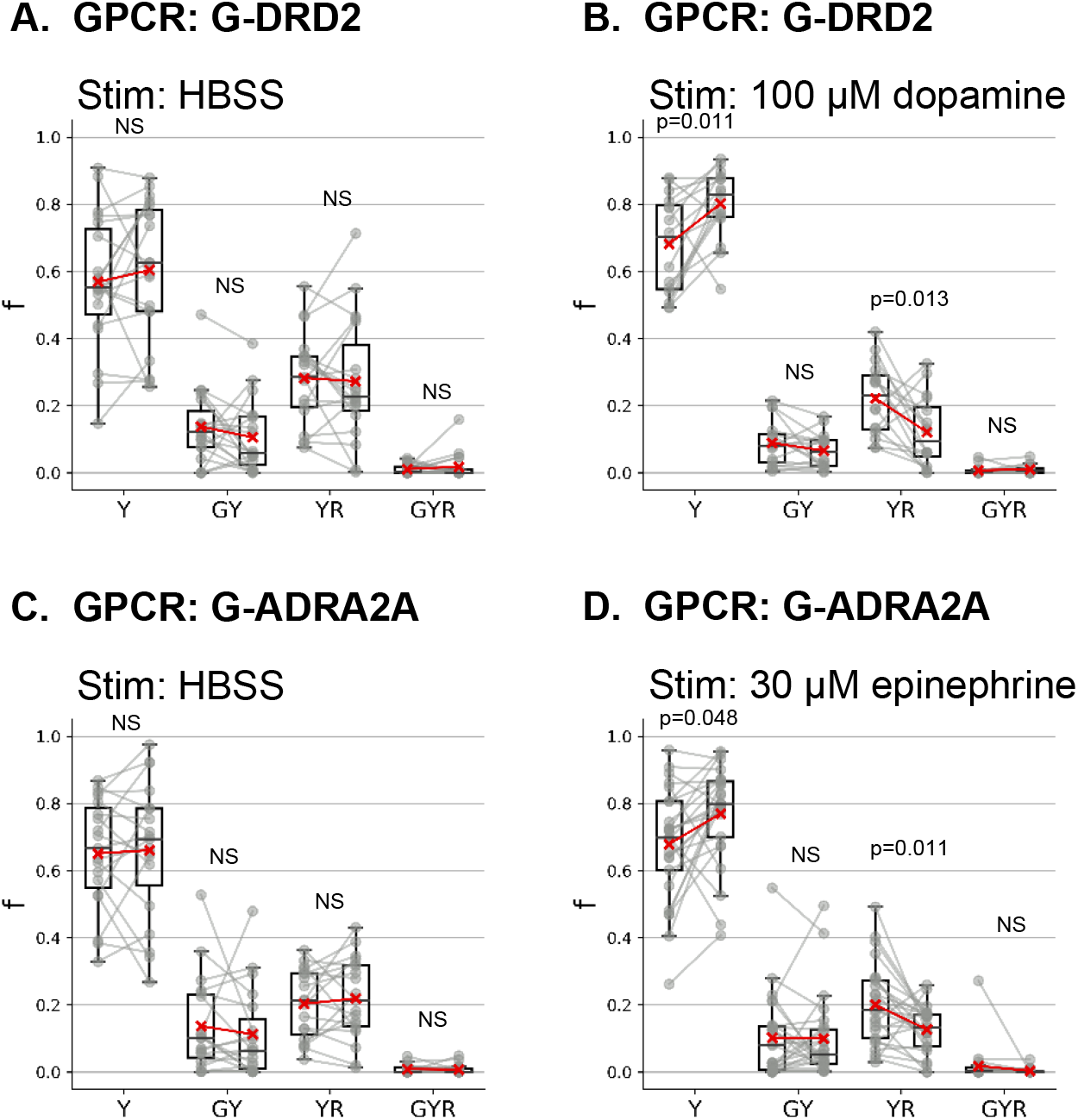
Fractional distribution of GNAI1-Y among oligomer states when coexpressed with R-GNB1, R-GNG2, and G-DRD2 or G-ADRA2A determined by covariance matrix analysis. For each state, the left column contains the pre-stimulus fraction, and the right column contains the post-stimulus fraction. A) G-DRD2 is the coexpressed GPCR with the G protein trimer components. Cells were stimulated with additional imaging buffer, HBSS (negative control). B) GPCR G-DRD2 stimulated with 100 μM dopamine. C) GPCR G-ADRA2A stimulated with HBSS (negative control). D) G-ADRA2A stimulated with 30 μM epinephrine. P-values are the results of two-sided paired t-tests.

The distributions of G-DRD2 and G-ADRA2A are dominated by noninteracting fractions consisting of ^~^90% of independent G chromophores (Fig. S3 A-B, Fig. S4 A-B). Apparent interactions with Y-GNAI1 and R-GNB1/R-GNG2 are comparable to background levels observed in control experiments (Fig. 2 A) and trimer (GYR) fractions were negligible.

R-GNB1/R-GNG2 also appears almost entirely as noninteracting with the other chromophores (>90%) (Fig. S3 C-D, Fig. S4 C-D). The apparent asymmetry between the fraction of Y participating in YR interactions and the fraction of R participating YR interactions comes from the relative expression levels of these two chromophores. Because GNAI1-Y expression is dictated by an internal ribosome entry site (Fig. S2), its expression is approximately three times lower than that of R-GNB1 and R-GNG2 (37).

We observe small relative changes in oligomer state distributions for G-GPCRs and R-GNB1/R-GNG2 in response to GPCR stimulation. The GR fraction of R chromophores increases after dopamine stimulation (Fig. S3 D), the GYR fraction of G chromophores decreases after epinephrine stimulation (Fig. S4 B), and the GYR fraction of R chromophores also decreases after epinephrine stimulation (Fig. S4 D). In each of these cases, the absolute oligomer fractions do not differ significantly from background levels (Fig. 2) and are unlikely to represent biologically relevant findings.

### Spectrally Resolved RICS Analysis of GPCR Stimulation Experiments

Datasets for FCMA of G protein activation were also processed using spectrally resolved RICS. The results from these analyses are in good agreement with those determined by FCMA. We observe negligible relative cross-correlations for GY and GR pairings (Fig. 4). Like FCMA, we observe YR relative cross-correlations that are sensitive to GPCR stimulation (Fig. 4 B, D).

**FIGURE 4.**
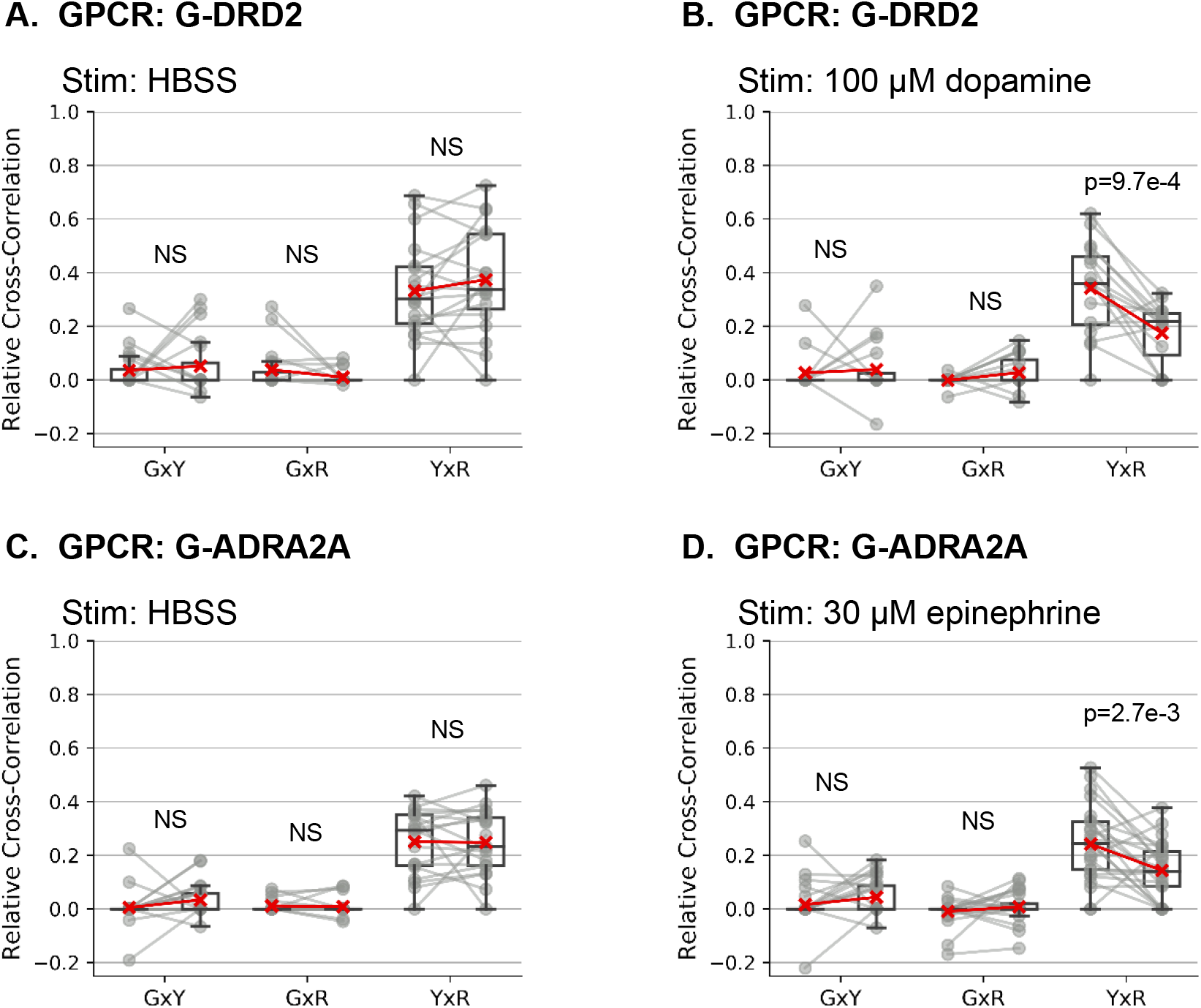
Relative cross-correlations determined by spectrally resolved RICS for three components of the G protein signaling cascade. For each chromophore pairing (i.e. GxY), the left column contains the pre-stimulus relative cross-correlations and the right column contains the post-stimulus cross-correlations. A) Relative cross-correlations for labeled GPCR G-DRD2 coexpressed with GNAI1-Y and R-GNB1/R-GNG2 stimulated with HBSS (negative control). B) Relative cross-correlations for G-DRD2, Y-GNAI1, and R-GNB1/R-GNG2 stimulated with 100 μM dopamine. C) Relative cross-correlations for G-ADRA2A, Y-GNAI1, and R-GNB1/R-GNG2 stimulated with HBSS (negative control). D) Relative cross-correlations for G-ADRA2A, Y-GNAI1, and R-GNB1/R-GNG2 stimulated with 30 μM epinephrine. P-values were determined by two-sided paired t-tests. NS denotes p>0.05.

### Diffusion Coefficients by Spectrally Resolved RICS

In contrast to FCMA, spectrally resolved RICS analysis also provides the apparent diffusion coefficient of each chromophore as an additional readout. While there are no statistically significant changes in the apparent diffusion coefficients for the GPCRs or G proteins in response to GPCR stimulation, there are distinct diffusivities among the components of the signaling cascade (Fig. 5). GPCRs, DRD2 and ADRA2A, are the least diffusive, averaging ^~^0.25 μm^2^/s. The R labelled GNB1/GNG2 component moves slightly faster than the receptors, averaging ^~^0.5 μm^2^/s. The Y labelled GNAI1 is the most diffusive, ^~^0.7 μm^2^/s, and shows a trend towards faster diffusion following stimulation of both GPCRs (p=0.071 with G-DRD2 and p=0.074 for G-ADRA2A) (Fig. 5 B, D).

**FIGURE 5.**
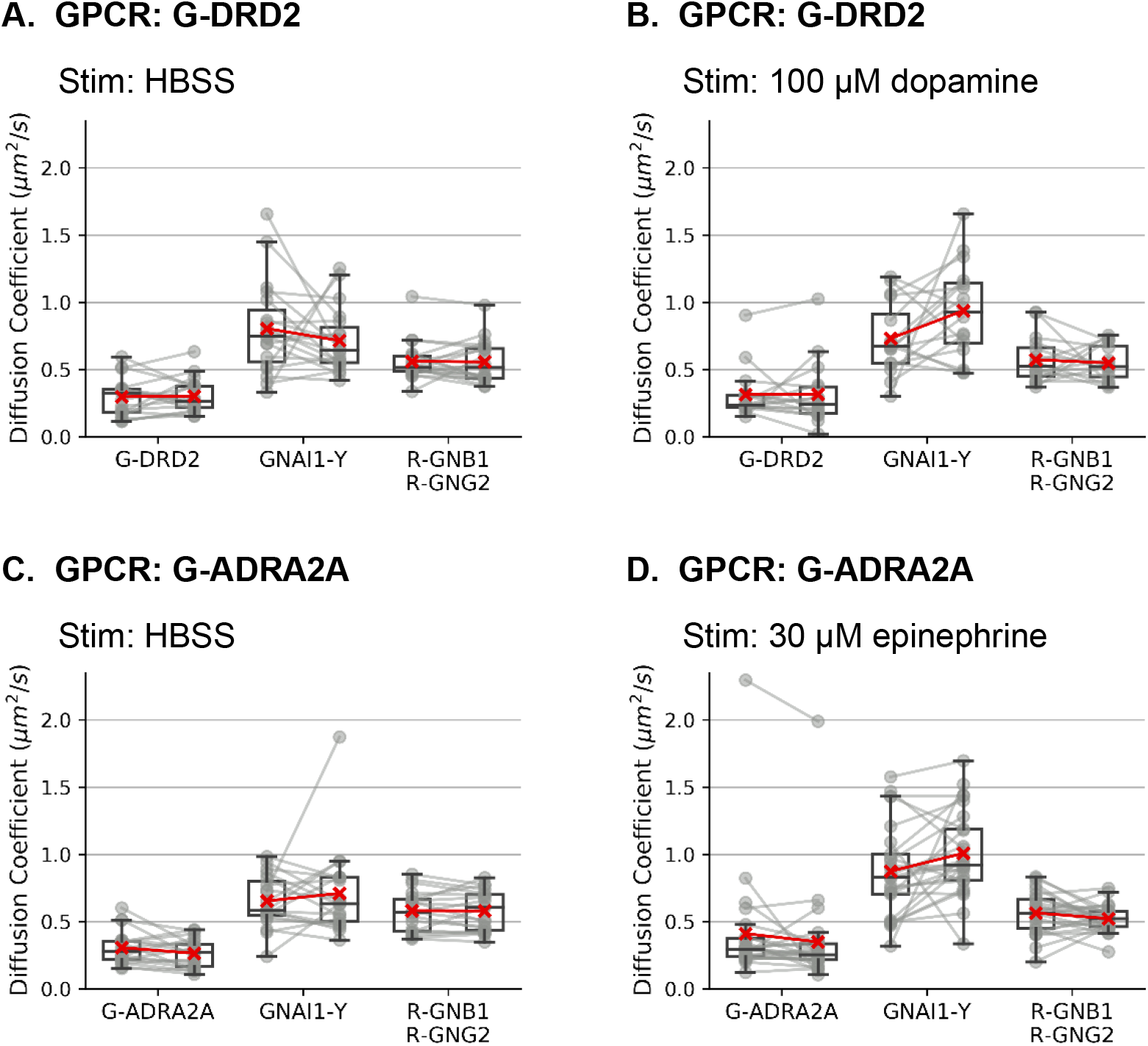
Diffusion coefficients for three components of GPCR signaling cascade determined from autocorrelation functions of spectrally resolved RICS. For each component, the left column contains the pre-stimulus diffusion coefficient, and the right column contains the post-stimulus diffusion coefficient. A) G-DRD2, GNAI1-Y, and R-GNB1/R-GNG2 stimulated with HBSS (negative control). B) G-DRD2, GNAI1-Y, and R-GNB1/R-GNG2 stimulated with 100 μM dopamine. C) G-ADRA2A, GNAI1-Y, and R-GNB1/R-GNG2 stimulated with HBSS (negative control). D) G-ADRA2A, GNAI1-Y, and R-GNB1/R-GNG2 simulated with 30 μM epinephrine. No changes between pre- and post-stimulation diffusion coefficients were found to be statistically significant (p>0.05, determined by two-sided paired t-test).

## Discussion

We have demonstrated FCMA as a fluorescence fluctuation analysis tool suitable for multicolor imaging experiments in live cells. In this analysis, heteromeric combinations leave unique fingerprints on the covariance matrices calculated from spectrally resolved images (Fig. 1). The relative contributions of different oligomer states can be determined from fitting model functions and the resulting information tells us how chromophores are distributed across these states (Fig. 2).

FCMA is complementary to recent developments in spectrally resolved image correlation spectroscopy. FCMA and spectrally resolved RICS achieve many of the same goals. Both quantify the degree of interaction between two or more chromophores (Figs. 2–4). RICS, and other correlation function-based approaches, have the advantage of providing information about transport properties by outputting fitted diffusion coefficients (Fig. 5). However, as we show in this work, FCMA detects ternary interactions directly (Fig. 2 B), which offers a more robust and simpler computational procedure. Ternary complex detection has been achieved with triple correlation analysis (TRICS), but that relies on higher order correlation functions which greatly increase the signal-to-noise requirements for the data and computational complexity (17, 38). Additionally, visual inspection of covariance matrices allows for the straightforward observation of complex formation that has an intuitive connection to the emission spectra (Fig. 1, Fig. S1 C). In practice, both analyses can be implemented in parallel with the same fluorescence imaging data.

The FMCA approach allows for simultaneous measurements of the three major components of the canonical GPCR/G-protein signaling pathway directly with fluorescence in live cells (Fig. 3, Fig. S3–S4). These data are highly relevant to our mechanistic understanding of the signal propagation through GPCR/G protein pathways (23). The two predominant models of G protein activation are collisional coupling (26–28) and pre-assembled complexes (29–31). In collisional coupling models, GPCRs and G proteins have independent Brownian motions aside from their brief interactions when the GPCR is activated. Conversely, pre-assembly models posit that stable GPCR/G protein complexes are present with the components maintaining contact throughout the signaling processes. Biochemically, the distinction between collisional coupling and pre-assembly models arises from the affinities of GPCR/G protein interactions (32). In this work, we did not find interactions between GPCRs (DRD2 and ADRA2A) and G proteins (Gαi1/Gβ1/Gγ2) (Fig. 3–4, Fig. S3–S4) that were distinguishable from background (Fig. 2). These data suggest that these GPCR-G protein components interact through weak, transient associations below what is detectable with our current experimental sensitivity. Additionally, we observed that the GPCR and G protein components have distinct apparent diffusion coefficients suggesting that they are not constitutively coupled as a pre-assembly model would suggest (Fig. 5). Although these data are consistent with a collisional coupling mechanism of GPCR signaling, we are limited by the fidelity of the chromophores to radiative states (Fig. 2 B-E) (39). Recently, spectrally resolved FFS for four chromophores has been demonstrated in live cell experiments, opening the door for the expansion of the work shown here to include more components of the signaling cascade such as downstream effectors (17).

## Methods

### DNA Plasmids

Subcloning to generate plasmids used in these studies was performed using standard molecular biology techniques. Most experiments utilized In-Fusion cloning reagents from Takara Bio Inc or ig-Fusion cloning reagents from Intact Genomics to facilitate subcloning unless noted otherwise. Reagents were used following the manufacturers’ protocols. Primers to generate linearized vectors and inserts are listed in Table S1.

Monomeric and multimeric fluorescent controls were derived by modifying previously published plasmids designed to express CD86-EGFP (Addgene # 133858) and mApple-CD86-EGFP (Addgene #133860) at the plasma membrane of mammalian cells (8). Parent plasmids were linearized using inverse polymerase chain reactions to allow for insertions of fluorescent protein gene substitutes. Three fluorescent proteins were used as labels in this work: mEGFP, mEYFP(Q69K), and mCherry2 (39–42), abbreviated as G, Y, and R, respectively. Monomeric controls, CD86-G, CD86-Y, and CD86-R were produced by making swaps against EGFP in CD86-EGFP. The dimeric controls R-CD86-G and R-CD86-Y were produced by making dual swaps against mApple and EGFP in mApple-CD86-EGFP. To generate CD86-Y-G and R-CD86-Y-G, an additional fluorescent protein site was introduced in the C-terminal linker regions of CD86-G and R-CD86-G, respectively. mEGFP encoding inserts were amplified from G-DRD2 described below. mEYFP(Q69K) was available from previous work in our lab (43). An mCherry2 donor plasmid (#54563) was obtained from Addgene thanks to a generous donation by Michael Davidson.

A polycistronic construct to express G proteins GNAI1, GNB1, and GNG2 with the fluorescent protein labels used in control experiments was created by modifying a plasmid introduced by van Unen et al (37). The parent plasmid, GNB1-T2A-cpVenus-GNG2-IRES-GNAI1-mTurquoise2, was obtained from Addgene (#69623). Both proteins of the Gβ1γ2 functional unit were labeled with R. R-GNB1 was generated by swapping R against EGFP in EGFP-GNB1 described previously (Addgene #133856) using restriction sites AgeI and BsrGI (8). GNB1-T2A-cpVenus-GNG2-IRES-GNAI1-mTurquoise2 was linearized about GNB1 by digestion with NheI and SacI, and R-GNB1 was inserted. An N-terminal label of GNG2 was introduced by linearizing the parent plasmid about its cpVenus encoding region and inserting R. Y was swapped against mTurquoise2 in the parent plasmid in two steps. First Y was swapped against mTurquoise2 in the monocistronic version of the plasmid, GNAI1-mTurqouise2 (Addgene #69620), using restriction digestion at AgeI sites to linearize the parent plasmid about mTurquoise2 and create GNAI1-Y by insertion of Y against mTurquoise2. BssHII and XmaI restriction sites were used to insert Y flanked by partial fragments of GNAI1 into the target vector using T4 ligation. A schematic of the final construct, R-GNB1-T2A-R-GNG2-IRES-GNAI1-Y is shown in Figure S2.

G-DRD2 was created from EGFP-DRD2, from previous work (8, 44), by using site directed mutagenesis to introduce the A206K modification to EGFP (40). To create G-ADRA2A, G-DRD2 was linearized about its DRD2 encoding region. ADRA2A was amplified from ADRA2A-Tango, obtained from Addgene (#66216) as a gift from Bryan Roth (45), and inserted against the position previously occupied by DRD2. All plasmids and their sequences will be made available via Addgene (https://www.addgene.org/Dave_Piston/).

### Cell Culture and Transfection

HEK293 cells were cultured in 1:1 Dulbecco’s Modified Eagle’s Medium/F-12 Ham with Glutamax+ supplemented with 10% fetal bovine serum albumin, penicillin, and streptomycin. Cells were incubated at 37 C with 5% CO_2_. Cells with passage number between 25 and 35 were used. For imaging, cells were transfected by electroporation. For a typical experiment, 1.5 x 10^6^ cells were electroporated with 1-3 ug of plasmid DNA per construct in a 2 mm gap cuvette and seeded among three four-chamber dishes with 30 mm diameters so that cells were at 50-75% confluency at the time of imaging. Electroporation was performed with eight 150 V pulses lasting 100 μs separated by 500 ms intervals. Cells were imaged 12-36 hours after electroporation.

### Confocal microscopy

All imaging experiments were conducting with a Zeiss LSM 880 confocal laser scanning microscope using a 40x, 1.2 NA, C-Apochromat water immersion objective lens. 488 nm was used to excite G and Y, and 561 nm was used to excite R. Typical laser intensities measured after the objective lens were 450 nW for 488 nm and 1.3 μW for 561 nm. A 488/561 main beam splitter was used to separate excitation and emission. For imaging of the basal plasma membrane for FCMA and spectrally resolved RICS analysis, scanning was performed over 256×256 pixels spanning a 13.28×13.28 μm^2^ area, 50 nm/pixel, 16.48 us/pixel, and 45 frames per cell. The 32-bin Quasar spectral detector recorded photoelectron counts by either photon counting or lambda mode in spectral bins of 8.9 nm spanning 490-695 nm (Figure S2).

Before imaging, cells were washed twice, and the growth buffer was replaced with Hanks’ Balanced Salt Solution (HBSS). During imaging cells were kept at 37 C with 5% CO_2_ using an incubated stage. For GPCR stimulation experiments, the same cells were imaged before and after the addition of the stimulus with approximately one minute of incubation between acquisitions.

### Fluorescence Covariance Matrix Analysis

FCMA is an extension of multicolor fluorescence cumulant analysis applied to spectrally resolved imaging data (8, 34, 35). For spectrally resolved imaging data the first order cumulants, *κ*_[1]_(*i*), are equal to the average detection spectrum of the pixels within the region of interest, R. The second order cumulants, *κ*_[1,1]_(*i, j*), are equal to the covariance matrix for all pairs of channels, (*i, j*).

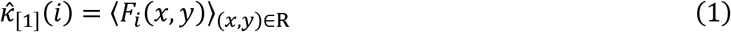

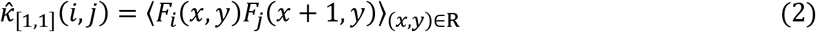

*F_i_*(*x, y*) denotes the fluorescence signal in photoelectrons at pixel (*x, y*) in channel *i*. To select regions of interest from images of basal plasma membranes we used a combination of polygonal selection and intensity thresholds of blurred images described previously (8, 46). Region of interest selection was performed on the spectrally unmixed images (Fig. S2 B-D) found for spectrally resolved RICS described below. To avoid complications from crosstalk in spectral detection we used a single pixel offset in the scanning axis (i.e. (*x* + 1)) discussed in depth in previous work (8).

Apparent number densities (*N_S_*) and molecular brightnesses (*ε_S_*) of species within the detection volume of the confocal microscope are related to the first and second order cumulants by:

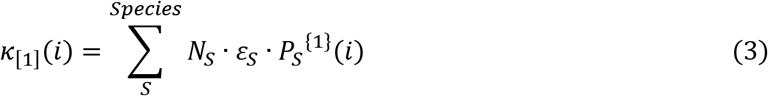

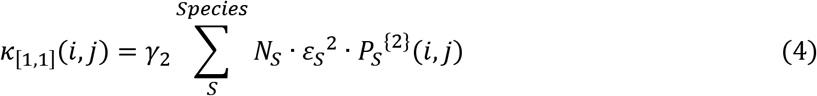

*γ*_2_ is a shape factor depending on the geometry of the detection volume. For this work we used *γ*_2_ = 0.5 corresponding to a two-dimensional Gaussian detection volume (47). *P_S_*^{1}^(*i*) for a species consisting of a single chromophore is equal to that chromophore’s detection spectrum normalized such that its sum is unity. *P_S_*^{2}^(*i, j*) is found from the outer product of *P_S_*^{1}^(*i*) with itself, *P_S_*^{2}^(*i, j*) = *P_S_*^{1}^(*i*) · *P_S_*^{1}^(*j*). When multiple chromophores are present in a species, *P_S_*^{1}^(*i*) is an average of its component parts weighted by the brightnesses of its constitutive chromophores (*C*):

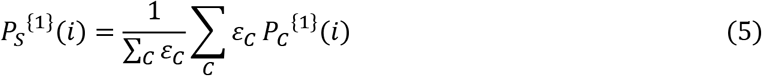

*P_C_*^{1}^(*i*) are determined in control experiments where a single chromophore is expressed. A consequence of Eqn. 5 is that species composed of unique combinations of chromophores are linearly independent.

For the three chromophore experiments performed in this work, we fit a seven species model to the detection spectra and covariance matrices. The number density was allowed to vary for each species and the molecular brightness was linked across species so that 10 variables were determined when fitting 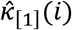 and 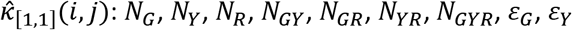, and *ε_R_*. Fitting was performed using Levenberg-Marquardt least squares minimization comparing experimentally determined 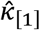 and 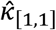 to theoretical values *κ*_[1]_ and *κ*_[1,1]_ (48, 49).

In this work, we focus on the distribution of number densities. Data are presented as fractional number densities. For example, for G containing species the fractional number density for species *x* is:

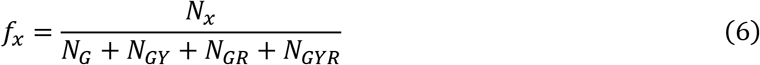
where *x* is *G, GY, GR*, or *GYR*.

### Spectrally Resolved Raster Image Correlation Spectroscopy

Spectrally resolved RICS was implemented following the approach established by Schrimpf et al (16) with some modifications introduced by Dunsing et al (17). For RICS analysis, the 23 channel raw images must first be decomposed into three-color images with each color corresponding to the fraction of that chromophore (Fig. S1). Regions of interest were specified using a combination of manual polygonal selection and intensity thresholding of blurred images (8). A temporal high pass filter was employed to exclude large fluctuations due to cellular movement and heterogeneity. The width of the high pass filter was three frames.

Spatial auto- and cross-correlation functions for each color and color pairing were calculated from binary masks using the arbitrary region algorithm described by Hendrix et al (46). Correlation functions were fit with a model function for two-dimensional diffusion:

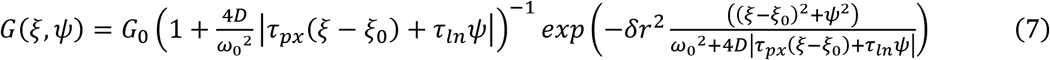

*ξ* and *ψ* are the lags along the scanning and scanning-orthogonal axes, respectively. *G*_0_ is the amplitude. *D* is the apparent diffusion coefficient. *ω*_0_ is the *e*^−2^ radius of the two-dimensional Gaussian detection area. *τ_px_* and *τ_ln_* are the pixel and line times, respectively. *δr* is the pixel size. *ξ*_0_ is the pixel lag along the scanning axis that produces the greatest fitted amplitude, *G*_0_.

We used the technique employed by Dunsing et al (17) to identify cross-correlation functions that did not have significant amplitudes. During fitting the initial *ξ*_0_ value was set to eight and if the fitted *ξ*_0_ was greater than three this was treated as having a nonsignificant correlation amplitude so that *G*_0_ = 0 for that experiment. This approach has the effect of ignoring spurious peaks at large pixel lags where the SNR is poor. To fit autocorrelation functions *ξ*_0_ was set to zero.

For the three chromophore experiments described in this work, spectrally resolved RICS generates six correlation functions, three autocorrelation functions (GG, YY, RR) and three cross-correlation functions (GY, GR, YR). Relative cross-correlations are calculated to quantify heteromeric interactions:

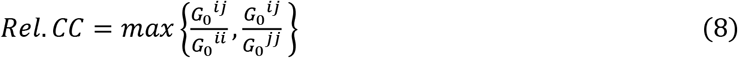

*i* and *j* specify the chromophore/channel.

As quality control measures, only data with SNRs greater than three in all autocorrelation functions and G/Y mean fluorescence ratios bounded by [1/10, 10] were included. SNRs were calculated using the method described by Schrimpf et al (16).

### Plotting and Statistics

Data are presented as swarm plots overlaid on box plots. The boxes indicate interquartile ranges. The centerlines indicate medians. Red exes indicate means. Whiskers are 1.5 times the interquartile ranges. Unless indicated otherwise, statistical tests were two-sided paired t-tests implemented with SciPy (50).

### Data and Code Availability

Analyses were performed using a series of IPython notebooks that are available at https://github.com/d-foust/fcma. Source data are available upon request.

## Author contributions

DJF and DWP planned the research. DJF performed the experiments, completed the analysis, and wrote the manuscript draft, which was edited by both authors.

## Acknowledgements

DJF received support from National Institutes of Health training grants T32EB14855 and T32DK108742. This research was supported by NIH grants R01DK123301 and R01DK115972 to DWP and by the Washington University Center for Cellular Imaging supported in part by the Diabetes Research Center (P30DK020579).

---

**FIGURE S1.**
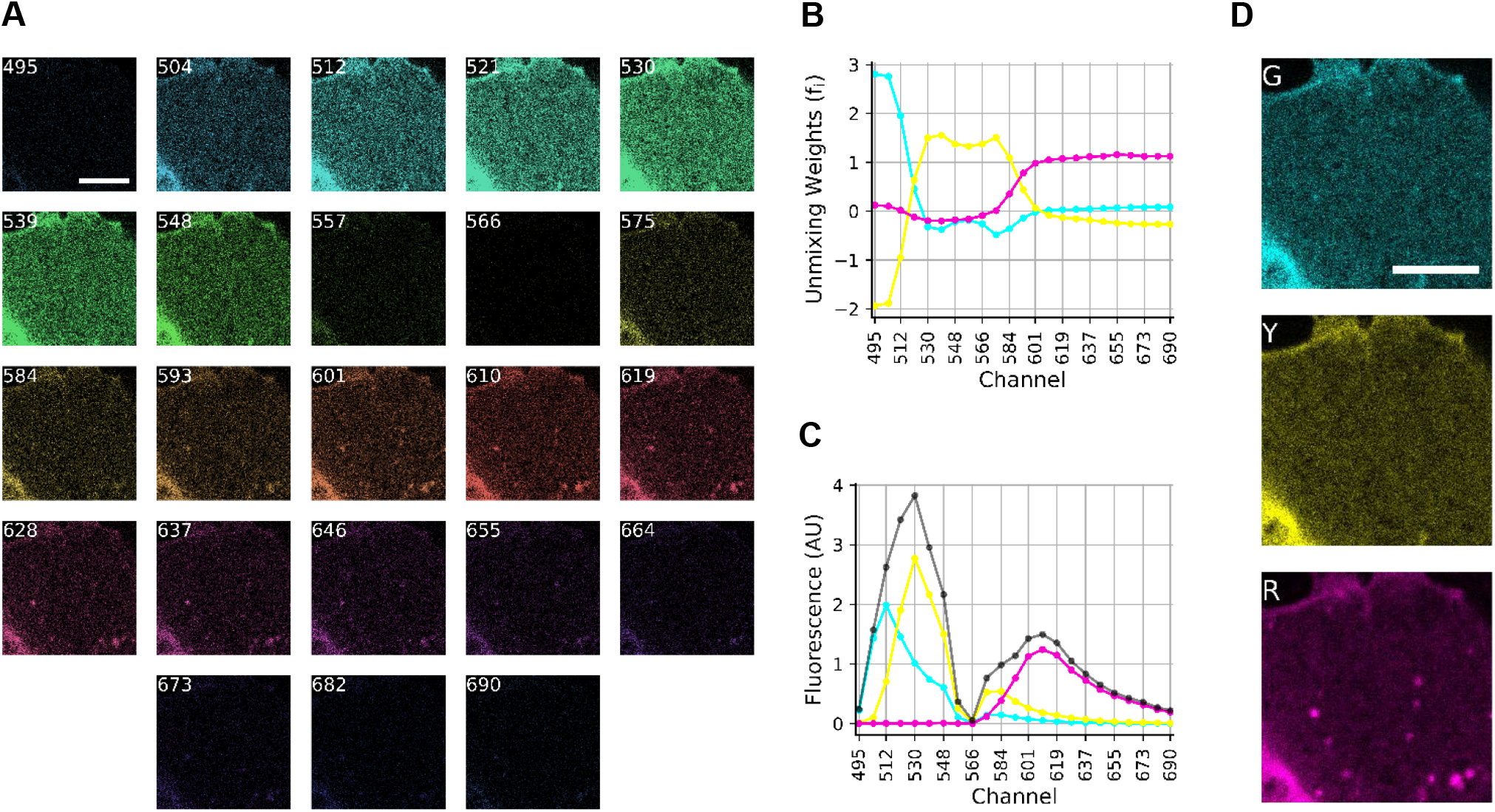
Spectral detection and unmixing for covariance matrix analysis and spectrally resolved RICS analysis. A) 23 channel image of the basal plasma membrane of a HEK 293 cell expressing R-CD86-Y-G. Labels denote the midpoint wavelength for each channel. B) Weights for spectral unmixing. Cyan, yellow, and magenta lines indicate weights for mEGFP, mEYFP(Q69K), and mCherry2, respectively. C) Detection spectra for image in panel A. The grey line is the composite detection spectrum. Cyan, yellow, and magenta lines represent contributions from mEGFP, mEYFP(Q69K) and mCherry2, respectively. D) Spectrally unmixed image corresponding to mEGFP, mEYFP(Q69K), and mCherry2. All scale bars are 5 μm.

**FIGURE S2.**
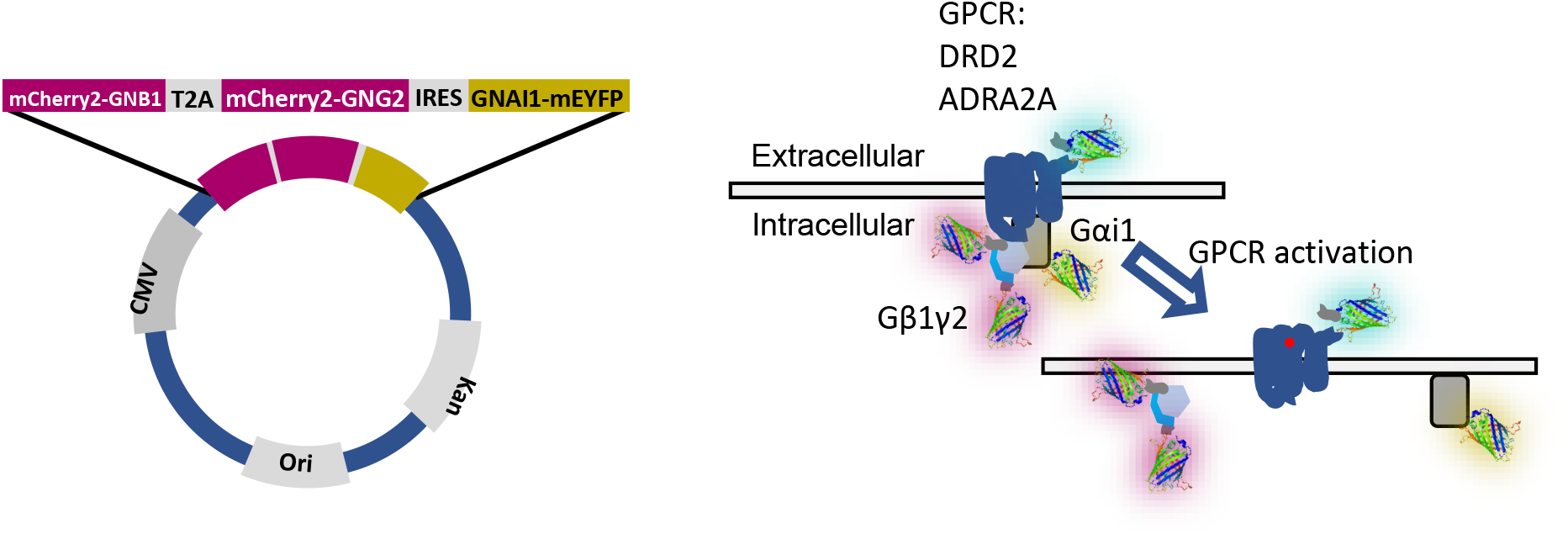
Left: Schematic of DNA plasmid used to express three components of the G protein heterotrimer. GNB1 and GNG2 are encoded with mCherry2 tags on their N-termini and separated by a T2A self-cleaving peptide. GNAI1 expression occurs under the control of an internal ribosome entry site and carries an mEYFP(Q69K) tag in the αb-αc loop of GNAI1. Right: Schematic of canonical G protein activation with pre-coupling. A ligand activated GPCR, labeled with mEGFP in this work, catalyzes the dissociation of the G protein complex into Gαi1 and Gβ1γ2 components.

**FIGURE S3.**
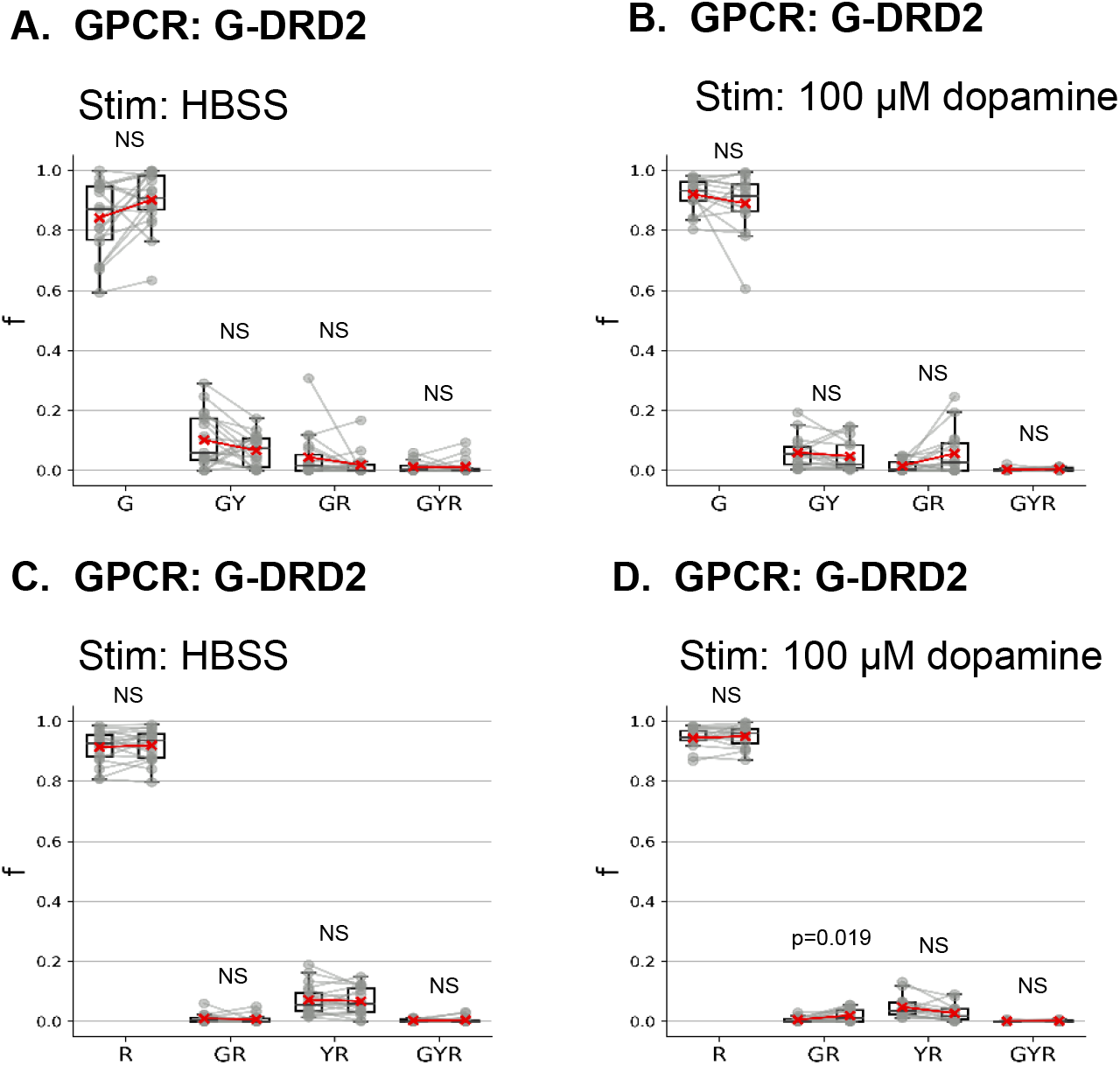
Fractional distributions of G-DRD2 and R-GNB1/R-GNG2 among oligomer states determined by covariance matrix analysis (corresponds with data in Figure 3 A-B). For each state, the left column contains the pre-stimulus fraction, and the right column contains the post-stimulus fraction. A) Fractional oligomer distribution of G-DRD2 stimulated with HBSS (negative control). B) Distribution of G-DRD2 stimulated with 100 μM dopamine. C) Distribution of R-GNB1/R-GNG2 stimulated with HBSS (negative control). D) Distribution of R-GNB1/R-GNG2 stimulated with 100 μM dopamine. P-values are the results of two-sided paired t-tests. NS denotes p>0.05.

**FIGURE S4.**
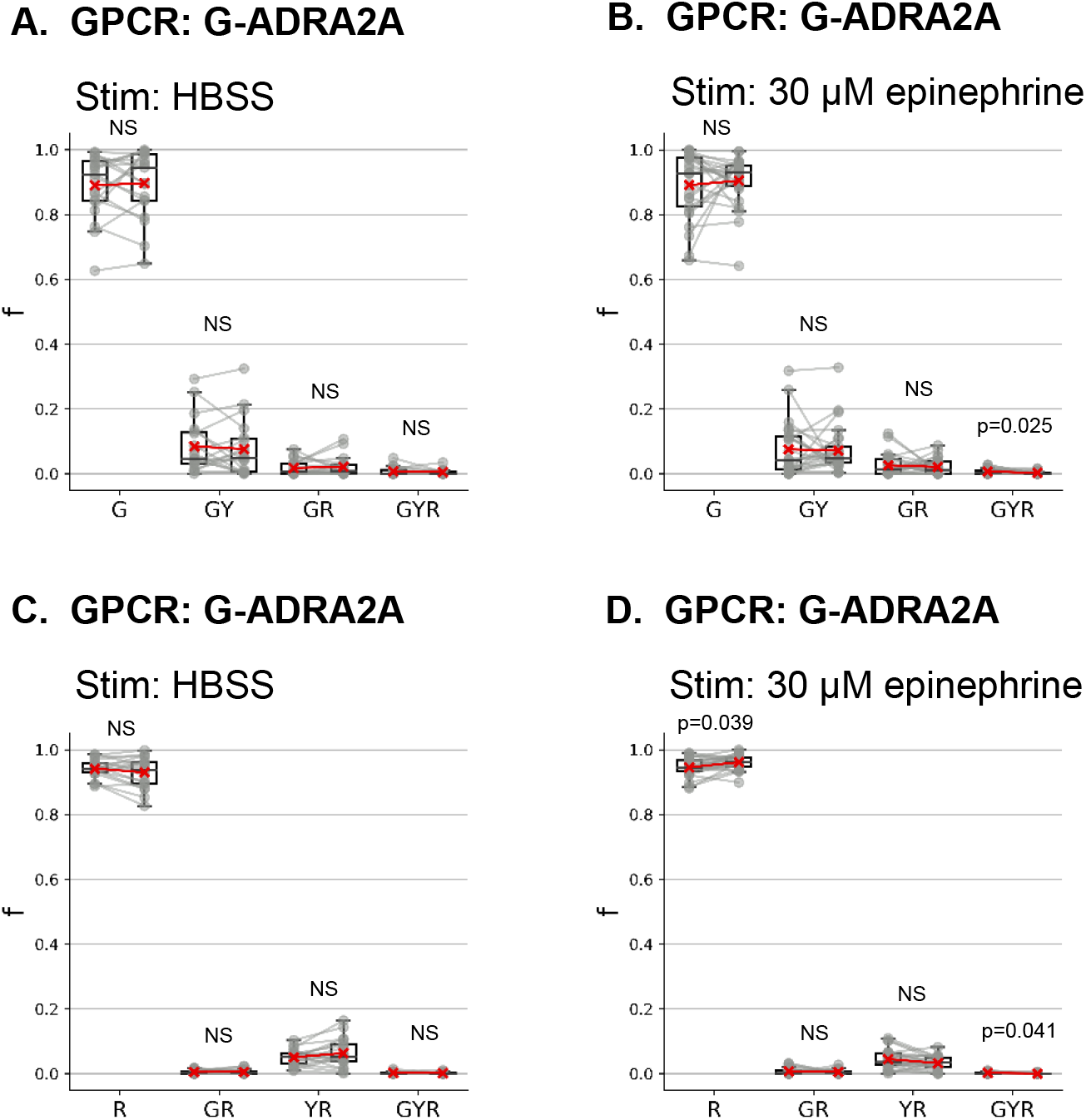
Fractional distributions of G-ADRA2A and R-GNB1/R-GNG2 among oligomer states determined by covariance matrix analysis (corresponds with data in Figure 3 C-D). For each state, the left column contains the pre-stimulus fraction, and the right column contains the post-stimulus fraction. A) Fractional oligomer distribution of G-ADRA2A stimulated with HBSS (negative control). B) Distribution of G-ADRA2A stimulated with 30 μM epinephrine. C) Distribution of R-GNB1/R-GNG2 stimulated with HBSS (negative control). D) Distribution of R-GNB1/R-GNG2 stimulated with 30 μM epinephrine. P-values are the results of two-sided paired t-tests. NS denotes p>0.05.

**Table S1.**
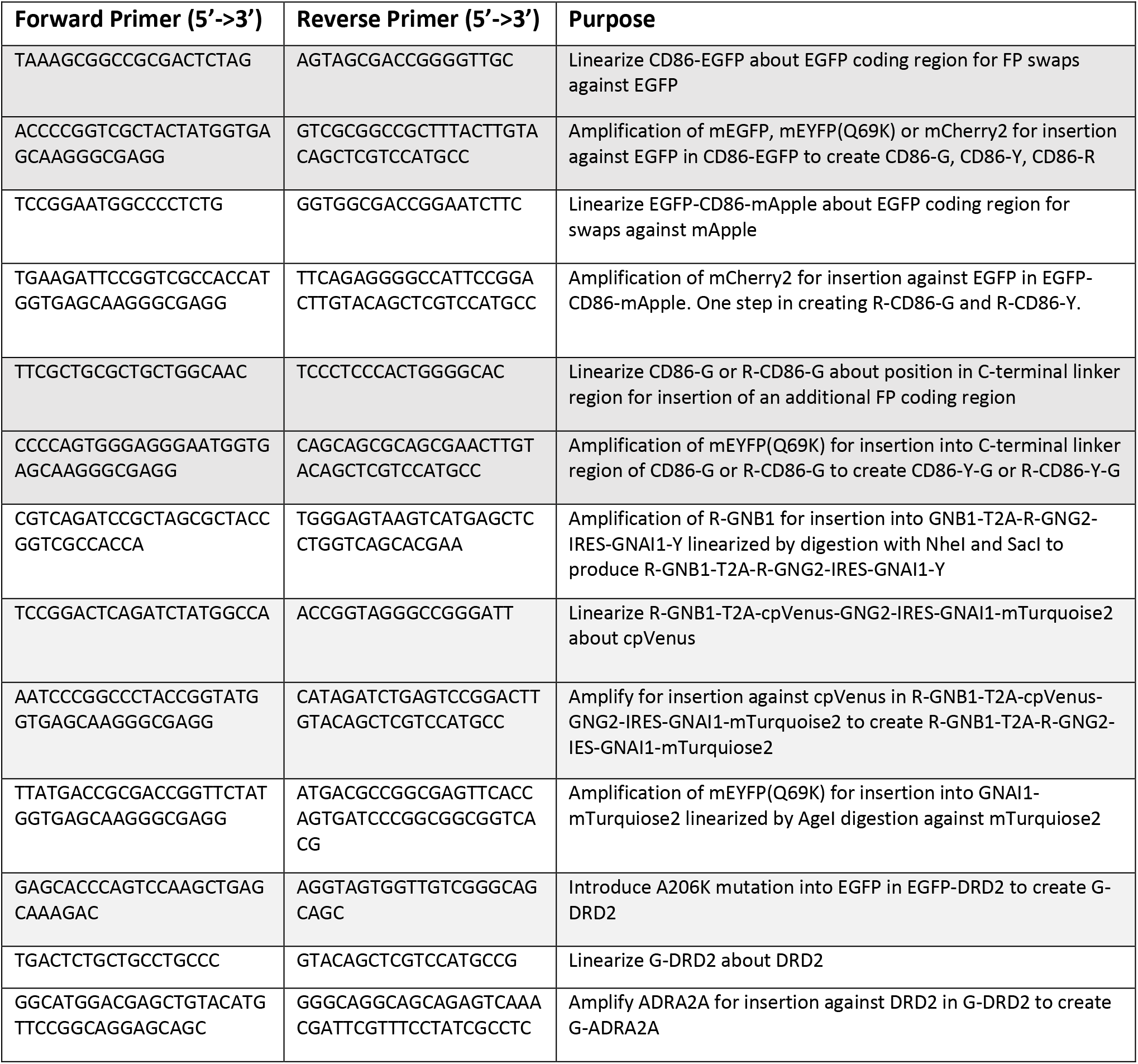
Primers for subcloning experiments.

